# The influence of birthweight, socioeconomic status, and adult health on brain volumes during aging

**DOI:** 10.1101/2023.09.21.558752

**Authors:** Christopher J McNeil, Tina Habota, Anca-Larisa Sandu, Gordon Waiter, Heather Whalley, Alison D. Murray

## Abstract

Preserved late-life brain volume is associated with resilience to dementia. We examined relationships between birthweight, socioeconomic status and adult health with late-life brain volumes. We hypothesised that early-life factors have direct and indirect effects on the aging brain. Neurotypical adults aged 61-67y underwent MRI and brain volumes measured. Birthweight, health and socioeconomic status were assessed by historical data or contemporary assessment. Measures of health and socioeconomic status were extracted using principal component analysis. Relationships between variables were explored by linear regression and structural equation models. Birthweight (β=0.095, p=0.017, n=280) and childhood socioeconomic status (β=0.091, p=0.033, n=280) were directly associated with late-life brain volume. Childhood socioeconomic status was associated with additional increase in grey matter volume (β=0.04, p=0.047, n=280). Better adult health was linked to increased late-life brain volume (β=0.15, p=0.003, n=280). Birthweight and childhood socioeconomic status are associated with whole and regional brain volume through direct and indirect mechanisms. Optimal fetal development, good adult health and reduced poverty, may prevent brain atrophy and decrease dementia risk in late-life.

## 1 Introduction

The health of the aging brain is the cumulative consequence of genetic and external factors experienced through life. These influence both brain development and growth in early-life, and deterioration and atrophy in later life. Greater brain volume is associated with resilience to pathology and better cognitive health in older people (Cox et al., 2016; van Loenhoud et al., 2018). In contrast, brain atrophy accompanies mild cognitive impairment (MCI) and Alzheimer’s dementia (Rana et al., 2016; Tabatabaei-Jafari et al., 2015). Aspects of the external environment, such as poverty and health, are highly inter-dependent and have complex interactions through life. Consequently, these factors could have direct effects or indirect effects on the brain. In addition, factors may not affect the brain, and merely co-vary with an influential factor.

Therefore, access to data regarding life-course factors that could affect brain volumes are required to analyse and understand how they interact, and their direct and indirect effects on the brain. Being small for gestational age (SGA) at birth is a consequence of intrauterine growth restriction (IUGR). IUGR is caused by maternal factors including poor nutrition (Belkacemi et al., 2010), environmental pollutants, (Wang et al., 1997) and smoking (Horta et al., 1997). Many of these factors are consequences of a poor environment during the mother’s pregnancy, often caused by poverty. IUGR effects the metabolism and body of the offspring, including the brain. Premature SGA babies have reduced brain and grey matter volumes at 1y and reduced brain volumes at 15y and 20y compared with those average for gestation age (AGA) (Ostgård et al., 2014; Rogne et al., 2015). Birthweight is associated with cortical area, and this different persists throughout life and consequently has life-long effects on general cognitive ability (Walhovd et al., 2016). These effects can persist into old age, and it has been measured that in adults aged 71-74y, their birthweight correlated with grey matter and whole brain volume, possibly due to changes to intracranial volume (Wheater et al., 2021).

Birthweight can predict the risk of poor adult health, and adult health can, in turn, affect brain volume. As first described by Barker and colleagues, low birthweight increases the risk of hypertension and type II diabetes (Whincup et al., 2008) and insulin resistance (Barker, 1997), both which can cause poor brain health.

There is mixed evidence that birthweight is associated with parental socioeconomic status (SES). SES was assessed using parental education and income and found children from a low SES household had lower birthweight and increased age 7y BMI (Morgen et al., 2017). In contrast, others did not observe a difference in household SES of SGA neonates compared with controls (Ostgård et al., 2014).

Childhood poverty and disadvantage has profound effects on brain volumes throughout the life course. Hanson et al. found that in children aged 7-18y, childhood SES was positively correlated with amygdala and hippocampal volume (Hanson et al., 2015). Childhood adversity and lower status paternal occupational have been associated with reduced brain cortical thickness, grey matter, and hippocampal volume in middle and late-life (Staff et al., 2012). Lower childhood SES is a strong risk factor for lower adult SES and for poor adult health (Cohen et al., 2010; Raaum et al., 2007). Poor adult SES correlates with smoking prevalence (Hiscock et al., 2012), hypertension (Grotto et al., 2008), type 2 diabetes (Hsu et al., 2012) and obesity (H. P. Booth et al., 2017), all associated with brain pathology.

From mid to late-life a higher adult SES was associated with greater brain volume and less brain pathology (AF et al., 2008; Waldstein et al., 2017). Low adult SES is a risk factor for AD, a condition characterised by regional and global brain atrophy (Kivimäki et al., 2020).

The interaction between poor adult health and SES is well established, and their close association is probably bidirectional. Impaired health often reduces economic activity and increase the risk of poverty and, conversely, lower SES is a risk factor for poor health due to reduced access to healthcare, lower levels of education and reduced engagement with health advice (Lago et al., 2018). Hypertension (Lane et al., 2019a; Newby et al., 2021), obesity (West et al., 2020), inflammation (Walker et al., 2017) and diabetes (Bresser et al., 2010; Roberts et al., 2014) are all associated with reduced regional or total brain volume and these conditions in midlife affect the brain in later life.

There is strong evidence for both direct, and indirect, effects of early-life on the health on the aging brain. The interconnected nature of life-course socio-economic status, health, and the brain makes the availability of whole life data on these factors required to understand their relationships, and their influence on the brain in late-life. This cohort is unique in having data for birthweight and cSES, collected contemporaneously, and not relying on participant recollection, as well as current measured of SES, health and brain MRI. This allows us to interrogate the direct and indirect determinants of brain health using reliable, longitudinal and detailed measures of the life-course environment.

Using data from a community living, older, non-demented sample we investigate if life course health and SES factors are associated with brain volume and structure in a sample. We use dimension reduction; structural equation modelling and a whole life dataset to test for associations to determine if factors could directly or indirectly affect the late-life brain. We hypothesised that birth weight, childhood SES and adult health affect brain size and structure in late life through direct mechanisms.

## 2 Methods

### 2.1 Participants

Participants were born between 1950-56 in Aberdeen, Scotland. In 1962 the Aberdeen Child Development Survey (ACDS) permanently linked various childhood records, including questionnaires completed by parents or guardians, to their school records and identity.

Between 1999 and 2001, 12150 members of ACDS were traced and 7183 were recruited into the Aberdeen Children of the Nineteen Fifties (ACONF). The ACONF participants completed a postal questionnaire that included recollections of childhood experiences, and questions of current health and SES. This data was linked to that collected in the ACDS and information collected by the Aberdeen Maternal and Neonatal Database (AMND) (Batty et al., 2004).

313 participants of the ACONF cohort were recruited into the STratifying Resilience and Depression Longitudinally (STRADL) study between 2015 and 2017 where a brain MRI scan, questionnaire of current health and SES, and physical measurements were performed. 280 had complete brain imaging data and were entered into this study (Habota et al., 2021).

### 2.2 Ethical approval and consent

This study received ethical approval from the Scotland A Research Ethics Committee (REC reference number 14/55/0039) and local Research and Development offices. All participants provided written informed consent. The ACONF dataset is registered with the National Research Ethics Service (NRES) as a research database.

### 2.3 Health, SES and family data

Between 1999-2001, aged 43-51y, ACONF participants completed a postal questionnaire of health and socioeconomic status (Leon et al., 2006). The questionnaire with questions relating to childhood socioeconomic status at age 12y, including: the occupation of their father, their family and house size and structure, use of a family car and parental tobacco use. Their birthweight and duration of pregnancy was obtained from the AMND. Relating to the participants current socio-economic status, current occupation, income, family size and structure, educational achievement, marital or cohabiting status and their car availability was recorded. Relating to the participants health: smoking and alcohol consumption, presence of a diagnosis of diabetes or hypertension. The participants were also asked to use bathroom scales to measure bodyweight and were asked their height.

These data were then used to calculate various socio-economic and health variables. Father’s occupation was defined using the standard occupational classification of 1990 (Surveys., 1990). This classification was then further analysed by socio economic group, the registrar’s general social class and by Nuffield classification (Goldthorpe et al., 1987). Living density (persons per room) was calculated from family and house size. Corrected birthweight was calculated by using linear regression to estimate the effect of pregnancy duration on birthweight. All birthweights were then corrected for a putative 40-week pregnancy duration.

The participants educational qualifications were assigned an ascending index based on the following: No formal qualifications, 0; School leaving certificate, 1; Other unspecified qualifications, 2; CSE ungraded, 3; CSE grades 2-5, clerical or commercial qualifications, 4; ‘O’ Grade passe, CSE grade 1 School certificate or matric City and Guild Craft/Ordinary level, 5; Certificate of Sixth Year Studies Highers A-levels ONC/OND/BEC/TEC (not higher) City and Guilds Advanced / Final level, 6; Teaching qualification HNC/HND, BEC/TEC higher, BTEC (higher) City and Guilds Full Technological Certificate, 7; Degree or degree level qualification (including a higher degree). Cigarettes smoking history was calculated from smoking status (never, past, current), duration of smoking, and packs smoked per week.

### 2.4 MRI and extraction of volumetric data

Participants were imaged on a 3T Philips Achieva TX-series MRI system (Philips Healthcare, Best, Netherlands) with a 32 channel phased-array head coil (software version 5.1.7; gradients with maximum amplitude 80 mT/m and maximum slew rate 100 T/m/s). Structural brain images were acquired using a 3D T1-weighted fast gradient echo with magnetisation preparation.

T1-weighted images were analysed using Freesurfer version 6.0 and segmentation checked for errors. Total intracranial volume (TICV), whole brain, grey matter and total hippocampal volumes were extracted.

### 2.5 Clinical and health measurements

Systolic and diastolic blood pressure (BP) was taken by a qualified research nurse using an automated sphygmomanometer (Omicron). Participants were asked to sit quietly in a warm room for five minutes and their blood pressure was measured in duplicate and mean BP used. Grip strength of both hands was measured with an automated dynamometer (Patterson Medical Jamar) and mean grip strength calculated. Height and weight were measured using calibrated scales and BMI calculated. Participants were asked their current or past smoking history, alcohol intake, presence of hypertension, diabetes, or other unspecified illness. The General Health Questionnaire (GHQ) was used to assess self-perceived psychological and physical well-being (Goldberg, 1978).

### 2.6 Extraction of childhood and adult SES and adult health principle components

Principle component analysis was used to extract a general composite measure for life-course factors described by multiple variables. General factors were extracted for a participant (occ) and their father’s occupation (f_occ) and used the subsequent PCA analyses. Childhood and adult SES (cSES gf and aSES gf) and adult health (AH gf) were then performed. The first un-rotated principal component, which best describes the variance shared between tests, was extracted.. For each factor extracted, the contributing variables, their contribution to the factor and their correlation with the extracted factor are detailed in tables 2-4. Details of the PCA extraction of f_occ and occ, and scree plots for cSES gf, aSES df and AH gf PCA extraction are detailed in Supplementary material.

### 2.7 Statistical analysis

Multiple regression was performed using intracranial, whole brain, grey matter and total hippocampal volume as the dependent variables. All models included age at scan and sex as covariates. For each dependent variable a ‘childhood’ and ‘whole life’ model were constructed. For the childhood model birthweight and cSES gf were included, and factors significantly associated with the dependent variable were included in the ‘whole life’ model. The whole life model included AH gf and aSES gf. For whole brain volume models, TICV was added as a confounding factor. For regional brain volumes models whole brain volume was included. Brain, rather than intracranial volume was included in these models to analyse regional volume effects, correcting for effects of global brain atrophy, which was analysed in a separate model. Statistical analysis was performed using R core (Schwarzer, 2007).

Structural equation models were constructed for those variables that were found by multiple regression to be significantly associated with volumes. The hypothesised structure of the path diagram was determined using the temporal sequence of factors or their structural relationship in the brain. Non-directional relationships were connected using a correlation term. Structural equation models were constructed using AMOS version 27 (Arbuckle, 2014). A goodness of fit for each model was assessed by chi-square significance (p>0.05), the goodness of fit, incremental fit and comparative fit indices (all > 0.95) and the Root Mean Square Error of Approximation (RMSEA, <0.08). Models fits of the data were acceptable when assessed using these indices and criteria. Model fit indices detailed in the Supplementary materials (Table a2).

## 3 Results

### 3.1 Sample summary

A summary of demographic, MRI and clinical data are shown in Table 1. 23 did not complete the MRI examination due to claustrophobia or the presence of safety exclusion criteria for MRI with the final N=280. There was no significant difference in birth weight, cSES gf, aSES gf, sex or AH gf between those with and without imaging.

**Table 1.** Summary of demographic, physical variables, and MRI derived brain volumes in Male and female participants.

| Variable | F, N = 155<sup>1</sup> | M, N = 125<sup>1</sup> | p-value<sup>2</sup> |
| --- | --- | --- | --- |
| Birth Weight (kg) | 3.26 (0.46) | 3.33 (0.48) | 0.2 |
| Age (y) | 62.28 (1.60) | 62.58 (1.52) | 0.10 |
| TICV (ml) | 1217.70 (206.91) | 1483.49 (185.47) | <0.001 |
| Brain Vol (ml) | 1006.32 (84.40) | 1119.30 (92.10) | <0.001 |
| Grey Matter Vol (ml) | 547.02 (42.12) | 604.67 (41.16) | <0.001 |
| White Matter Volume (ml) | 431.99 (45.65) | 486.42 (53.55) | <0.001 |
| <sup>1</sup>Mean (SD) |  |  |  |
| <sup>2</sup>Two Sample t-test |  |  |  |

### 3.2 Data Dimension Reduction by PCA

Participant and paternal occupation details were graded using 4 methods, and paternal occupation added information from the ADAS reading survey. These variables were used to extract the first unrotated principal component factor composite occupation measure. The paternal occupation component (f_occ) accounted for 78% of the total variance. The participants occupation component (occ) accounted for 82% of the variance.

Table 2 details the variables used to extract the first principal component, cSES gf, that describes childhood SES cSES gf accounted for 24% of the total variance amongst the contributing variables. Seven variables describe family wealth or ‘class’, four relate to health, and nine variables relate to family structure (size and participants status within family). Wealth variables contribute 34.7%, family structure variables contribute 52.2% and health variables contribute 13.1% to cSES gf. cSES gf is positively correlated with family affluence, a smaller family size and fewer people per room. Table 3 details the variables used to extract the first principal component that describes adult socio-economic status (aSES gf). aSES gf accounted for 35% of the total variance. Three variables describe family wealth or ‘class’, two relate to educational achievement and 3 relate to family structure (size and structure of family). Wealth variables contribute 49.2%, family structure variables contribute 33.3% and educational achievement variables contribute 17.5% to aSES gf. aSES gf positively correlated with wealth, a larger family size and greater educational achievement.

**Table 2.** Summary of variables for PCA analysis of cSES gf. Summary: Structure, family size and status amongst siblings; Health, health environment; Affluence, wealth or social status of parents. Contribution, contribution of variable to the 1^st^ principle component of AH. N=280.

| **Variable** | **Type** | **Mean (range)** | **Contribution (%)** |
| --- | --- | --- | --- |
| Family size in 1962 | Structure | 3.00 (1.00, 9.00) | 14.6 |
| Status index child | Structure | 2.02 (1.00, 9.00) | 12.3 |
| Number of older brothers, sisters | Structure | 1.07 (0.00, 8.00) | 11.9 |
| Pregnancy number | Health | 2.23 (1.00, 9.00) | 10.3 |
| Occupation density (person/room) | Affluence | 1.30 (0.33, 2.67) | 8.0 |
| Father's occupation PC | Affluence | 0.00 (-3.40, 4.06) | 7.5 |
| Number of children when 12y | Structure | 2.66 (1.00, 8.00) | 6.8 |
| Social class husband-survey 1962 | Affluence | 4.29 (1.00, 8.00) | 6.5 |
| Husband's social class birth of | Affluence | 4.43 (1.00, 8.00) | 6.5 |
| Number of adults when 12y | Structure | 2.26 (1.00, 5.00) | 3.6 |
| Family car at 12y | Affluence | 1.52 (1.00, 2.00) | 3.2 |
| Wife's pre-marital occupation | Affluence | 4.11 (1.00, 9.00) | 3.1 |
| Number of younger siblings | Structure | 1.08 (0.00, 10.0) | 1.5 |
| Number of living rooms at 12 years | Structure | 1.48 (1.00, 5.00) | 1.4 |
| Wife's height | Health | 15.8 (1.0, 29.0) | 1.3 |
| Age at delivery-complete years | Health | 27.2 (18.0, 41.0) | 0.8 |
| Parents smoking | Health | 2.00 (1.00, 4.00) | 0.7 |
| Brought up by both parents | Structure | 0.94 (0.00, 1.00) | 0.1 |
| Number of bedrooms at 12 years | Structure | 2.60 (0.00, 6.00) | 0.0 |
| Father's employment type | Affluence | 3.01 (1.00, 9.00) | 0.0 |

**Table 3.** Summary of variables for PCA analysis of aSES gf. Summary; Structure, family size; Wealth, income or wealth, Educational Achievement, level of academic qualification achieved; Contribution, contribution of variable to the 1^st^ principle component of aSES gf. N=280.

| **Variable** | **Type** | **Mean (range)** | **Contribution (%)** |
| --- | --- | --- | --- |
| Participants Occupation PC | Wealth | 0.00 (-4.84, 2.17) | 20.6 |
| Qualification Score | Educational Achievement | 5.82 (0.00, 8.00) | 17.5 |
| Income per week | Wealth | 5.26 (0.00, 8.00) | 14.9 |
| Number of children living with | Structure | 1.01 (0.00, 4.00) | 14.8 |
| Number of cars | Wealth | 1.52 (0.00, 2.00) | 13.7 |
| Living with husband, partner | Structure | 1.47 (1.00, 4.00) | 10.3 |
| Number of offspring | Structure | 1.77 (0.00, 8.00) | 8.2 |

Table 4 details variables used to extract the first principal component for the adult health factor AH gf. AH gf accounted for 19% of the total variance. Nine variables were collected at ages 43-51y and nine aged 59y to 67y. Fourteen variables were self-reported and four were directly measured. Six variables were classified as ‘health behaviour’ seven as ‘metabolic related’ and five as ‘general health’. General health variables contribute 18.6%, health behaviour variables contribute 62.4% and metabolic health variables contribute 19% to AH gf. AH gf was negatively correlated with smoking, alcohol intake and measures of metabolic disease risk.

**Table 4.** Summary of variables for PCA analysis of AH. Summary: Behaviour, health related behaviour; General, general indices of health; Metabolic, relating to obesity, hypertension or diabetes. AU = Arbitrary Units. U = 10 ml of ethanol. Contribution, contribution of variable to the 1 principle component of AH. N=280.

| Variable | Type | Mean (range) | Contribution (%) |
| --- | --- | --- | --- |
| Health last 12m (AU) | General | 1.80 (1.00, 4.00) | 16.1 |
| Current illness | General | 0.13 (0.00, 1.00) | 13.5 |
| Hypertension (59-67y) | Metabolic | 0.28 (0.00, 1.00) | 9.7 |
| BMI | Metabolic | 26.9 (17.6, 48.4) | 9.3 |
| GHQ score (59-67y) | General | 15 (6, 61) | 8.3 |
| Cigarettes smoked | Behaviour | 60,886 (0, 723,195) | 8.1 |
| Hypertension (43-51y) | Metabolic | 0.12 (0.00, 1.00) | 7.9 |
| Systolic BP (mmHG) (59-67y) | Metabolic | 141 (76, 194) | 6.7 |
| Current illness (59-67y) | General | 0.71 (0.00, 1.00) | 6.4 |
| Diastolic BP (mmHG) (59-67y) | Metabolic | 83 (46, 118) | 6.1 |
| Smoker? (AU (59-67y) | Behaviour | 1.2 (0.0, 30.0) | 4.7 |
| Diabetes (43-51y) | Metabolic | 0.0107 (0.0000, 1.0000) | 2.9 |
| Alcohol intake (u/wk) (43-51y) | Behaviour | 4.6 (0.0, 30.0) | 0.4 |
| Alcohol intake (u/wk) (59-67y) | Behaviour | 8 (0, 60) | 0.1 |
| Grip strength (kg) (59-67y) | General | 28 (7, 55) | 0.0 |

### 3.3 Multiple regression and structural equation analysis

Table 5 shows the results of two multiple regression models that test the effect of childhood and adult factors on TICV. The models include the confounding factors of sex and age at MRI scan. In both models only sex significantly predicted TICV. Being male predicted a 0.5 SD (∼120 ml) larger intracranial volume.

**Table 5.** Regression results of the association between life course factors and TICV. Standardised coefficients with t-values. N=280.

|  | TICV |  |
| --- | --- | --- |
|  | Childhood Factors | Adult Factors |
| Sex | 0.556<br>t = 11.073 *** | 0.537<br>t = 10.464 *** |
| Age | -0.032<br>t = -0.641 |  |
| Birth Weight | 0.064<br>t = 1.280 |  |
| cSES gf | 0.031<br>t = 0.617 |  |
| Adult Health |  | -0.017<br>t = -0.305 |
| aSES gf |  | 0.100<br>t = 1.824 |
| R<sup>2</sup> | 0.317 | 0.319 |
| Adjusted R<sup>2</sup> | 0.307 | 0.312 |
| Residual Std. Error | 0.833 (df = 275) | 0.830 (df = 276) |
| F Statistic | 31.883 *** (df = 4; 275) | 43.126 *** (df = 3; 276) |

*Note:* \* p<0.05, \*\* p<0.01, \*\*\* p<0.001

Table 6 shows the effects of childhood and adult factors on total brain volume. Brain volume was not affected by age at scan. Testing childhood factors found that sex and TICV both predicted brain volume with being male predicting a 0.2 SD (21 ml) increased brain volume independent of the effect of increased TICV. For each SD increase TICV predicted a 0.6 SD increase in brain volume (1 ml increased TICV predicted a 0.47ml increased BV).

**Table 6.** Regression results of the association between life course factors and brain volume. Standardised coefficients with t-values. N=280.

|  | Brain Volume |  |
| --- | --- | --- |
|  | Childhood Factors | Adult Factors |
| Sex | 0.212<br>t = 4.429<sup>***</sup> | 0.226<br>t = 4.750<sup>***</sup> |
| Age | -0.068<br>t = -1.714 |  |
| Intracranial Volume | 0.591<br>t = 12.366<sup>***</sup> | 0.580<br>t = 12.3<sup>***</sup> |
| Birth Weight | 0.097<br>t = 2.448<sup>*</sup> | 0.094<br>t = 2.39<sup>*</sup> |
| cSES gf | 0.108<br>t = 2.722<sup>**</sup> | 0.091<br>t = 2.15<sup>*</sup> |
| Adult Health |  | 0.151<br>t = 3.67<sup>***</sup> |
| aSES gf |  | -0.040<br>t = -0.89 |
| R<sup>2</sup> | 0.573 | 0.589 |
| Adjusted R<sup>2</sup> | 0.565 | 0.580 |
| Residual Std. Error | 0.659 (df = 274) | 0.648 (df = 273) |
| F Statistic | 73.566<sup>***</sup> (df = 5; 274) | 65.9<sup>***</sup> (df = 6; 273) |

Note: \* p<0.05, \*\* p<0.01, \*\*\* p<0.001

Birth weight predicted BV, with each SD increase predicting an increased brain volume of 0.1 SD. This equates to 22 ml increased BV per 1 kg birth weight increase (2% of BV). An increased cSES gf score predicts greater brain volume. The magnitude of effect of birthweight and cSES gf was approximately equivalent. The effect of birthweight and cSES gf remained significant when adult factors were included. aSES gf did not predict brain volume when other factors were considered. Brain volume was significantly associated with adult health, with each SD increase in adult health resulting in an 0.17 SD increase in BV (equivalent to 18 ml BV). A Structural equation model of these data is shown in Figure 1. This analysis supported the findings made with multiple regression, showing that birth weight correlated with aSES gf, cSES gf was correlated with AH gf and aSES gf, and there was a strong correlation between aSES gf and AH gf.

**Figure 1.**
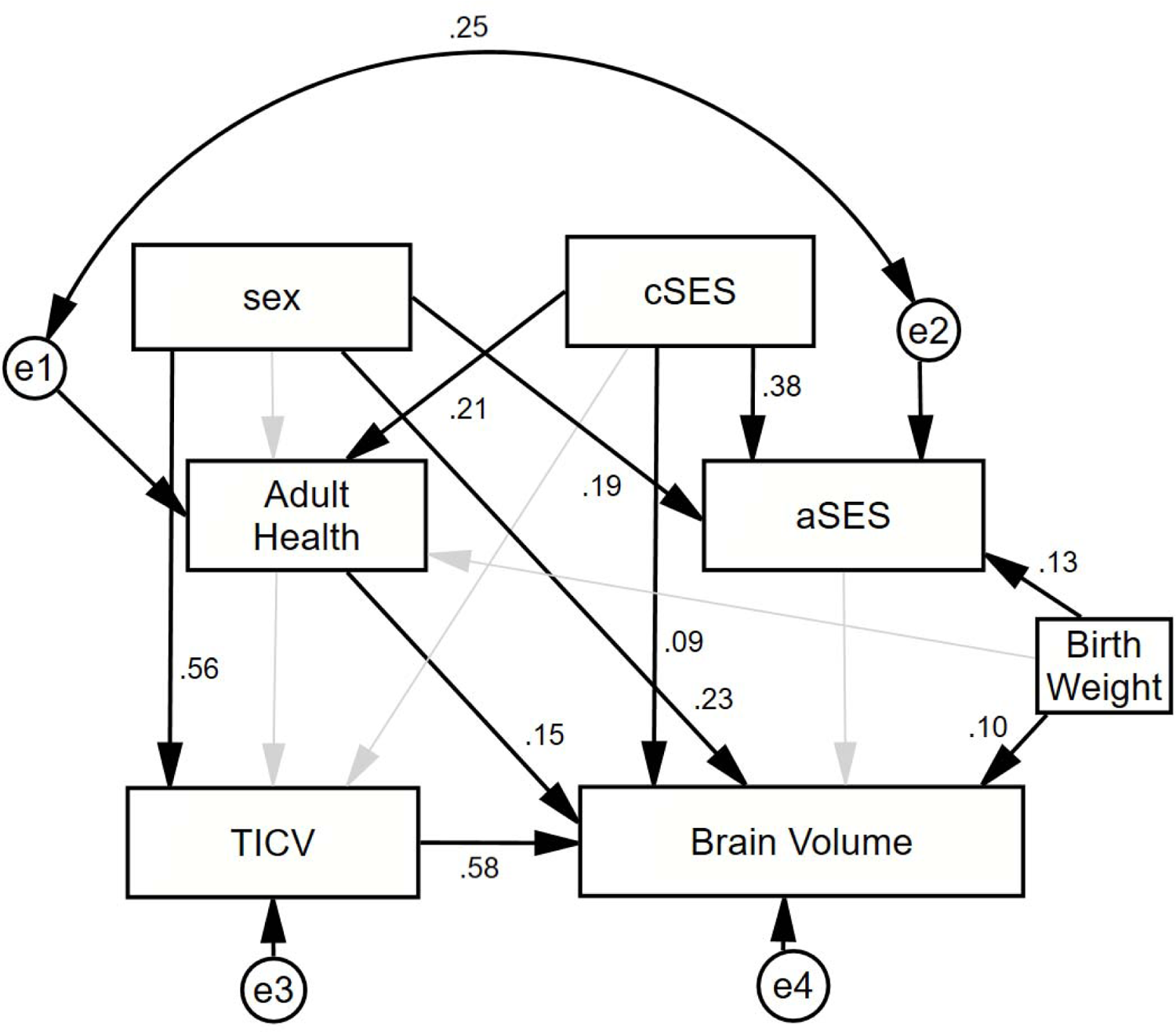
Path diagram of the structural equation model of the relationships between life course factors and brain volume. Non-significant standardised coefficients are not shown. Non-significant relationships represented by a light grey arrow.

Table 7 shows the results of multiple regression analysis of factors influencing grey matter and hippocampal volume. All models included brain volume as a covariate, so any effects detected were in addition to their previously detected effects on brain volume. Male participants had a significantly greater volume of grey matter. cSES gf significantly increased grey matter volume, with an increase of 2ml per SD cSES gf. No adult factors measured significantly influenced grey matter volume. Hippocampal volume was associated with brain volume, but not with the other factors included in the model. Figure 2 shows the structural equation model of determinants of grey matter volume.

**Figure 2.**
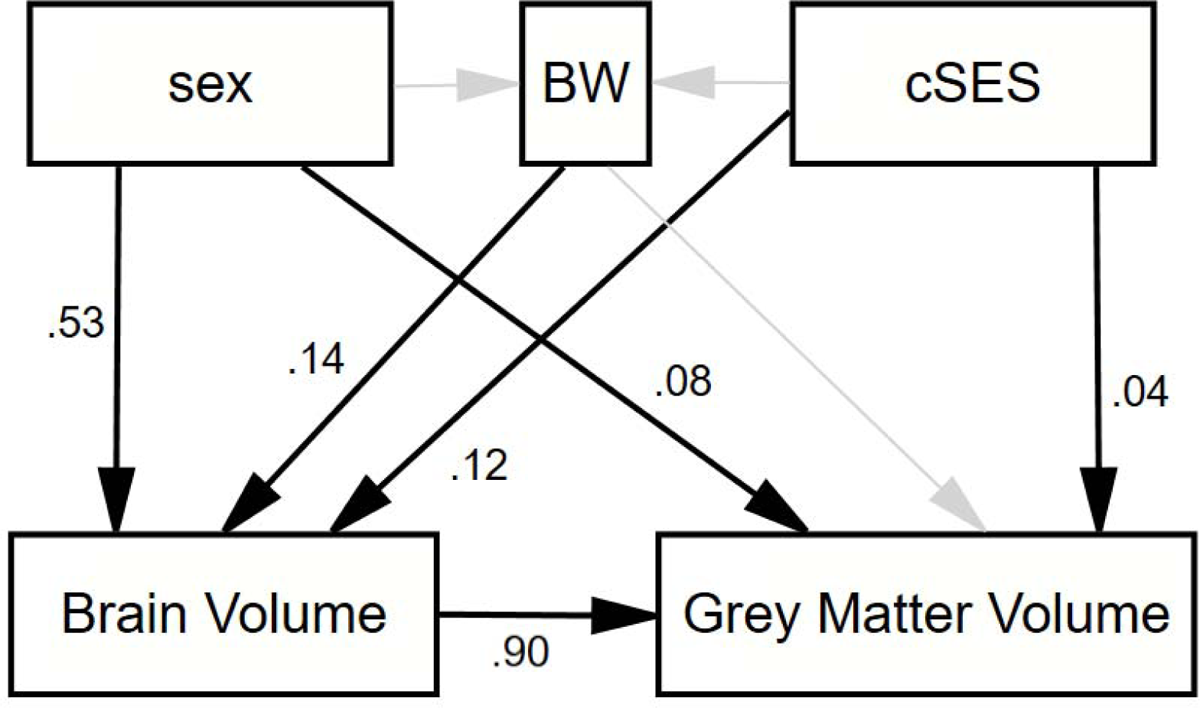
Path diagram of the structural equation model of the relationships between life course factors and grey matter volume. Non-significant standardised coefficients are not shown. Non-significant relationships represented by a light grey arrow.

**Table 7.** Regression results of the association between life course factors, grey matter and hippocampal volumes. Standardised coefficients with t-values. N=280.

|  | Grey Matter |  | Hippocampi |  |
| --- | --- | --- | --- | --- |
|  | Childhood Factors | Adult Factors | Childhood Factors | Adult Factors |
| Sex | 0.082<br>t = 3.776*** | 0.080<br>t = 3.617*** | 0.006<br>t = 0.117 | 0.007<br>t = 0.137 |
| Age | -0.001<br>t = -0.070 |  | -0.019<br>t = -0.428 |  |
| Brain Volume | 0.897<br>t = 40.561*** | 0.896<br>t = 40.249*** | 0.696<br>t = 13.026*** | 0.665<br>t = 12.403*** |
| Birth Weight | 0.027<br>t = 1.479 |  | -0.084<br>t = -1.905 |  |
| cSES gf | 0.036<br>t = 1.998* | 0.024<br>t = 1.197 | 0.024<br>t = 0.549 |  |
| Adult Health |  | 0.018<br>t = 0.908 |  | 0.043<br>t = 0.913 |
| aSES gf |  | 0.026<br>t = 1.274 |  | 0.049<br>t = 1.042 |
| R<sup>2</sup> | 0.911 | 0.911 | 0.480 | 0.478 |
| Adjusted R<sup>2</sup> | 0.909 | 0.910 | 0.471 | 0.470 |
| Residual Std. Error | 0.301 (df = 274) | 0.301 (df = 274) | 0.728 (df = 274) | 0.728 (df = 275) |
| F Statistic | 559.545*** (df = 5; 274) | 561.862*** (df = 5; 274) | 50.628*** (df = 5; 274) | 62.886*** (df = 4; 275) |
| Note: |  |  | * p<0.05, ** p<0.01, *** p<0.001 |  |

## 4 Discussion

We demonstrate that birth weight, childhood SES and adult health are associated with late-life brain volume through direct and indirect mechanisms. In addition, those who had a poor childhood SES had lower grey matter volume. Better adult health was correlated with greater brain volume. Intracranial and hippocampal volume were not associated with birth weight, childhood SES, adult SES or adult health. Birthweight, SES and health acted directly and indirectly, throughout the life-course on late-life regional and whole brain volumes.

### 4.1 Birthweight and brain growth and development

We found that birthweight was associated with whole brain volume during late-life, independent of a mechanism acting through SES or adult health. Brain volumes are affected by their environment before birth. In utero, fetal growth is sensitive to nutrition, environmental pollutants, smoking tobacco, and maternal stress hormones that can affect birthweight (Belkacemi et al., 2010; Diego et al., 2006; Wang et al., 1997). These factors can also affect pre- and postnatal brain development (Catena et al., 2019; Deoni et al., 2018; Ekblad et al., 2015; Teicher et al., 2016), and it is probable that the factors that affect fetal growth also affect brain growth and development. It is possible that birthweight is a proxy measure of factors that have altered prenatal brain development and growth.

### 4.2 Childhood SES and brain growth and development

Childhood SES had a direct association with late-life brain volumes. During early-life, the brain undergoes periods of development with simultaneous development and pruning of synapses accompanying hypertrophy of brain tissue (Stiles & Jernigan, n.d.). This period of growth can last until 20y and is influenced throughout by SES. Although SES can be defined in several ways (Shavers, 2007) it is commonly agreed that it is influenced by family size and wealth. Reduced SES can affect nutrition, health care access, education, and environment. Disruption of these factors during this crucial period may influence and have permanent consequences on brain structure and metabolism.

### 4.3 Consequences of differences in childhood SES and birth weight

The association we observe of birthweight and childhood SES on late-life brain volumes may directly operate by two, non-mutually exclusive, mechanisms. The first possible mechanism is that their influence can change the structure of the brain, resulting in greater initial volume and altered proportions of tissue. These changes may persist five decades later, and cause differences seen in brain volumes. It has been seen that education is a source of cognitive reserve (Staff et al., 2004), and many other childhood factors can also increase brain volume and provide a mechanism for providing reserve.

The second mechanism is via accelerated atropy, with evidence that early-life experiences can alter vulnerability to neurodegeneration. Early-life factors, including nutrition, alter late-life epigenetic methylation and are associated with late-life brain volumes (Gabbianelli & Damiani, 2018; Lorgen-Ritchie et al., 2021; Staff et al., 2012). Low birthweight and childhood stress also have profound effects on adult hormonal regulation, and the function of the immune system (Bilbo & Schwarz, 2009; Schlotz & Phillips, 2009). Immune system dysfunction and exposure to stress hormones are associated with brain conditions such as AD (Blasko & Grubeck-Loebenstein, 2003; Hatzinger et al., 1995). Birthweight and childhood SES may act on the brain through changes in epigenetic programming of metabolism or adult hormonal and immune system function.

### 4.4 Birth weight, childhood SES and hippocampal volume

We did not detect an effect of early-life factors on hippocampal volume. Evidence for the association of childhood SES with hippocampal volume in late-life is mixed. Studies have found that childhood SES was correlated with childhood, but not adult, hippocampal volume (Yu et al., 2018). It has been shown that childhood mistreatment, but not childhood SES, affected adult hippocampal volume (Lawson et al., 2017). Here, the cohort we studied may not have yet reached the age where evidence of increased atrophy of hippocampus caused by reduced childhood SES had either started or was detectable by MRI. When modelling hippocampal volume, we included brain volume as a confounding variable and hypothesised a change specific to the hippocampi. Other findings of hippocampal volume changes may have partially been a consequence of whole brain changes that were not included in the analysis.

### 4.5 Relationships between life-course health, SES, and the brain

We found that adult health was associated with late-life brain volumes, potentially by directly promoting brain atrophy or alternatively because adult health could share a common underlying cause with poor brain health. Hypertension, diabetes, and obesity are associated with reduced brain volume (T. Booth et al., 2013; Hsu et al., 2012; Lane et al., 2019b). A major contributor to AH df is smoking. Smoking tobacco is a major risk for the other health variables and is associated with brain atrophy (Durazzo et al., 2012). Considering smoking is crucial when investigating brain health in older populations, as it may be a primary, underlying cause of poor general and brain health. Birthweight was associated with adult SES, potentially by reducing educational achievement (Murthy et al., 2019) a key contributing variable for aSES gf, and an influence on other constituent variables of aSES gf, including wealth (Fergusson et al., 2008).

Adult SES did not affect brain structure directly but was strongly correlated with adult health. It is thought that brain volume is particularly vulnerable to SES during early-life, and during adult life it is health, rather than SES, that affects the brain. It is likely that changes in SES from childhood to adulthood through social mobility may benefit the brain through better adult health.

### 4.6 How important are these changes relative atrophy during dementia?

We can use these findings to predict the consequences of increased birthweight and childhood SES on cognitive reserve. The cumulative effect of a one SD increase in birthweight and cSES gf is 3-4% of brain volume. During normal aging, approximately 0.3% loss per year was constant through the life course in healthy participants (Scahill et al., 2003). A meta-analysis reports a loss of 1% per year during mild cognitive impairment (MCI) (Tabatabaei-Jafari et al., 2015). It has been proposed that brain volume can act as a source of physical cognitive reserve and resist the effects of neuro-pathology. These data suggest modest differences in birthweight and childhood SES could markedly influence late-life cognitive reserve.

### 4.7 Strengths

We used multiple measures of SES and adult health, and therefore avoided the limitations of using a single or small number of variables that may not be individually meaningful. In this way, we attempt to model the wider experience of an individual during their life. We used independent measures of adult health and SES, allowing us to separate the effects of these correlated and interdependent factors on brain volumes.

We included data collected from throughout the life course using both retrospective questionnaires and contemporaneous measurement, allowing longitudinal analysis of these data. These data do not have the inaccuracies and potential bias of data based on retrospective recollection of potentially emotive data about childhood environment and family SES. We exploited this longitudinal dataset by using structural equation modelling, allowing causal hypotheses to be examined.

We included intracranial or whole brain volumes as a confounding factor when modelling factors that affect volumes. Therefore, these findings for the whole brain, or its components, are specific for the region, and are not simply due to global volume changes.

### 4.8 Weaknesses

Using PCA analysis to create SES and AH composite scores resulted in AH, cSES and aSES gf that did not contain 60-80% of the variance within the original data. Using this method allows the large number of potentially important factors to be analysed using parsimonious and inclusive methods, but it is not possible to find specific mechanisms, and could miss effects caused by a single factor. It is not possible to prove causal relationships between variables in this long-term human cohort study, therefore it is possible that unmeasured factors may influence the relationships we observed. For example, altered fetal growth or childhood SES could affect late-life brain volumes through intermediate unmeasured mechanisms.

### 4.9 Conclusion

During late-life, the volume of the whole brain, and its grey matter, are independently associated with birthweight and childhood socioeconomic status from 6 decades earlier. Importantly, it seems these associations are direct and are not entirely explained by changes to adult health, which is itself associated with late-life brain volumes. These findings suggest that early-life factors can directly affect late-life brain volume, either through changes in initial volumes during development or accelerated atrophy in later life. We also found that adult health is an important determinant of late-life brain volumes. Early-life experiences may directly alter resilience to neuropathologies including Alzheimer’s Dementia.

## Acknowledgements

The imaging assessment was supported by The Wellcome Trust grant 104036/Z/14/Z to CM, AM, HCW and Andrew McIntosh.

## 5 Disclosure Statement

Declarations of interest: none.

## Notes

### Competing Interest Statement

The authors have declared no competing interest.

