## Supplementary Material for "The influence of birthweight, socioeconomic status, and adult health on brain volumes during aging"

### a. Supplementary Materials

#### a.1 PCA of Fathers Occupation

Table a‑1.

| **Characteristic** | **N = 280***^1^* |
| --- | --- |
| q1f_soc | 594 (110, 990) |
| q1f_se91 | 7.65 (1.20, 16.00) |
| q1f_rg91 | 3.13 (1.00, 5.00) |
| q1f_gd91 | 4.92 (1.00, 7.20) |
| rs031 | 41 (1, 91) |
| *^1^* Mean (Range) | |

Figure a‑1

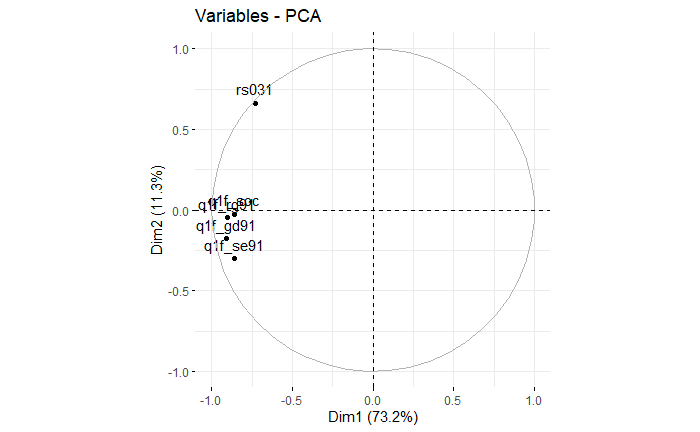

#### a.2 PCA of participants occupation

Figure a‑2 general factor

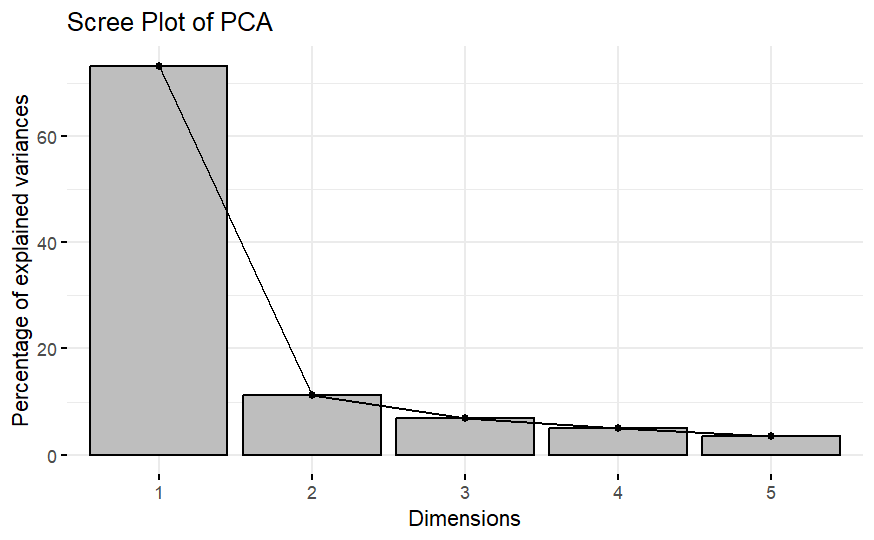

#### a.3 PCA of participants occupation

| **Characteristic** | **N = 280***^1^* |
| --- | --- |
| Standard Occupational Classification | 391 (102, 990) |
| Socio-Economic Group 1991 based on SOC | 5.22 (1.10, 14.00) |
| Registrar-Generals Social Class 1991 based on SOC | 2.56 (1.00, 5.00) |
| Goldthorpe 1990 | 2.89 (1.00, 7.10) |
| *^1^* Mean (Range) | |

#### a.4 PCA of Childhood Socio-Economic Status (cSES)

Figure a‑3. Scree plot of Childhood Socioeconomic Status general factor (cSEC gf) principle component analysis

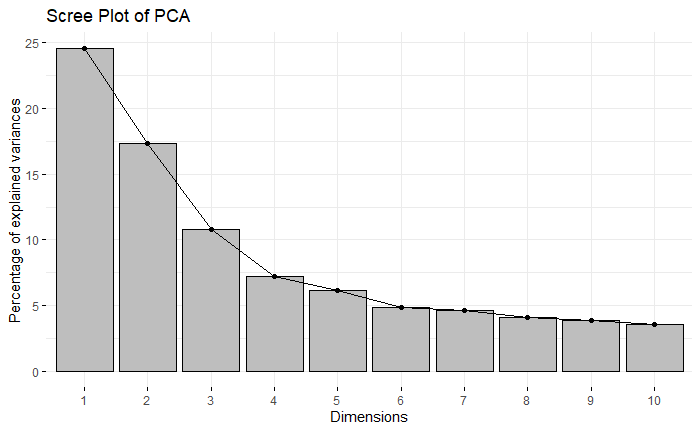

#### a.5 Adult Socioeconomic Status (aSEC gf) principle component analysis

Figure a‑4. Scree plot of Adult Socioeconomic Status general factor (aSEC gf) principle component analysis

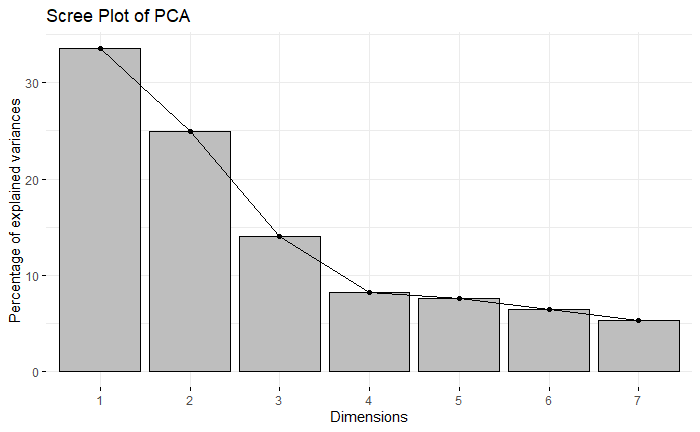

Figure a‑5. Scree plot of adult health general factor (AH gf) principle component analysis

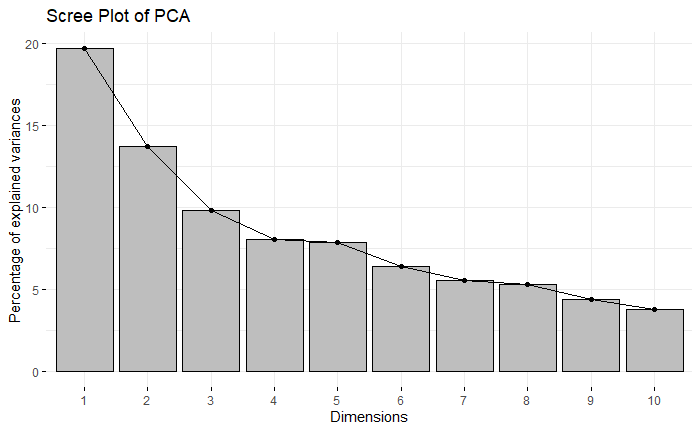

#### a.6 Structural Equation Model fit indices

Table a‑2. Fit indices of SEM analysis detailed in Figures 1 and 2.

|  | SEM Model | |
| --- | --- | --- |
| Model Fit Metric | Figure 1 | Figure 2 |
| Chi Squared | 0.56 | 0.835 |
| Goodness of Fit (GFI) | 0.99 | 1 |
| Comparative Fit index (CFI) | 1 | 1 |
| Root Mean Square Error of Approximation (RMSEA) | 0.074 | 0 |

#### a.7 Added variable plots of significant relationships of tables 5 to 7

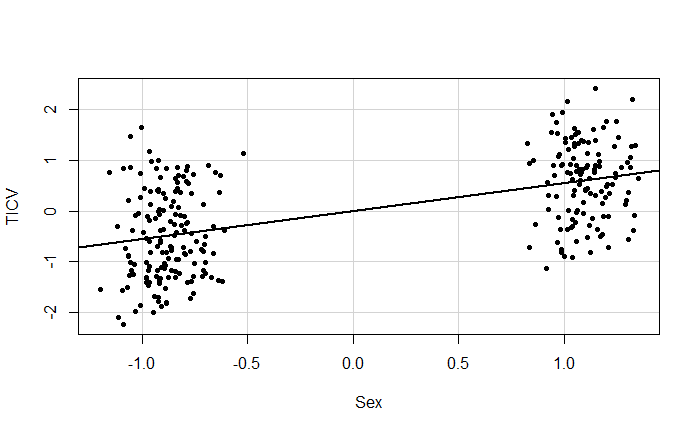

Figure a‑6. Added variable plot of the effect of sex on TICV, from Table 5, childhood factors. Sex; -1 = Female, +1 = Male. TICV = standard deviations. This plot demonstrated the effect of sex on TICV, correcting for the other factors in the multiple regression.

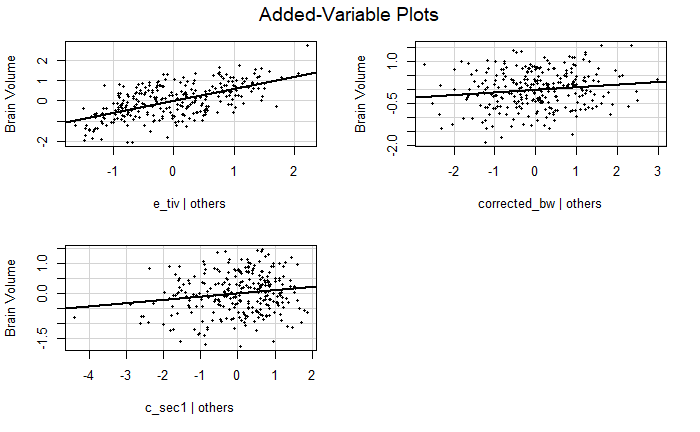

Figure a‑7. Added variable plots of the effect of TICV, BW and cSEC on BV, from the multiple regression detailed in Table 6: childhood factors. All values are standard deviations. These plots demonstrate the effect of the independent variable on BV, correcting for the other factors in the multiple regression.

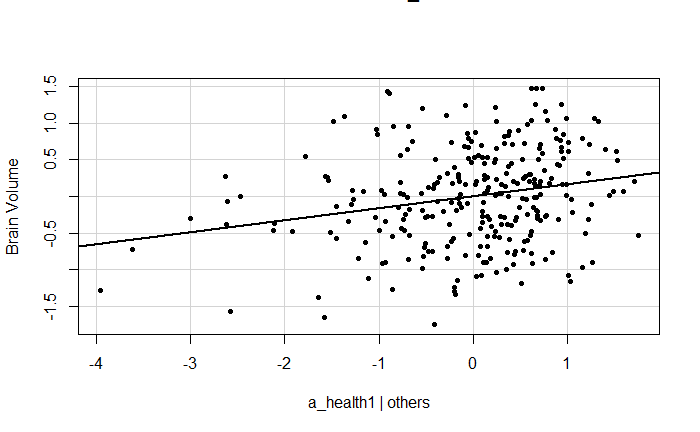

Figure a‑8. Added variable plot of the effect of Adult Health on BV, from the multiple regression detailed in Table 6: adult factors. All values are standard deviations. This plot demonstrates the effect of the independent variable on BV, correcting for the other factors in the multiple regression.

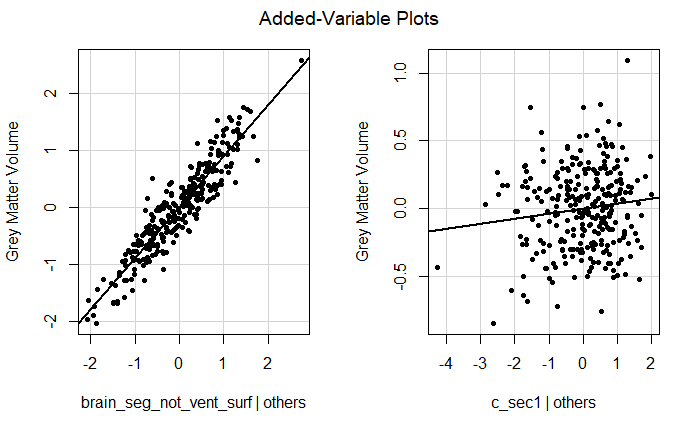

Figure a‑9. Added variable plot of the effect of BV and cSES on grey matter volume, from the multiple regression detailed in Table 7: childhood factors. All values are standard deviations. This plot demonstrates the effect of the independent variable on grey matter volume, correcting for the other factors in the multiple regression.

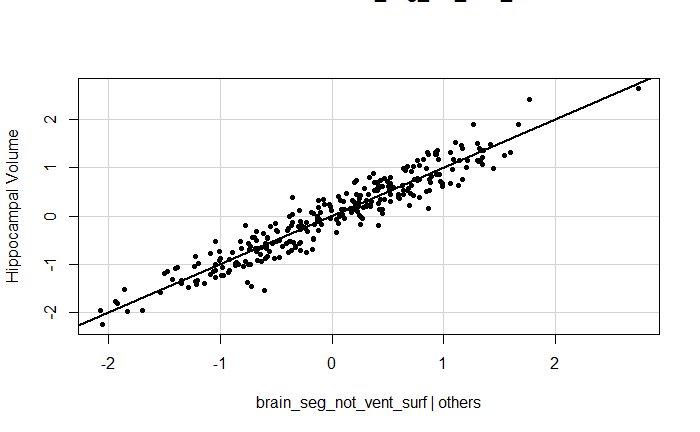

Figure a‑10. Added variable plot of the effect of BV on hippocampal volume, from the multiple regression detailed in Table 8: childhood factors. All values are standard deviations. This plot demonstrates the effect of the independent variable on hippocampal volume, correcting for the other factors in the multiple regression
